# Comparative genomics of canonical antifungal dsRNA target genes across fungal plant pathogens: conserved siRNA-length windows define the scope for broad-spectrum spray-induced gene silencing

**DOI:** 10.64898/2026.11.01.763432

**Authors:** Muhammad Wasif, Muhammad Ahsan Khan, Muhammad Abubakkar Azmat, Gábor Tarcali, Huma Saleem

**Affiliations:** Institute of Plant Protection, Faculty of Agricultural and Food Sciences and Environmental Management, University of Debrecen, Böszörményi út 138, 4032 Debrecen, Hungary; Department of Plant Breeding and Genetics, University of Agriculture, Faisalabad 38000, Punjab, Pakistan; Department of Plant Breeding and Genetics, University of Agriculture Faisalabad, Constituent College Burewala, Vehari 61010, Pakistan

**Keywords:** Spray-induced gene silencing, dsRNA biopesticide, Comparative genomics, Fungal plant pathogens, Broad-spectrum, siRNA, CYP51, β-tubulin, Chitin synthase

## Abstract

**Background:** Spray-induced gene silencing (SIGS), in which double-stranded RNA (dsRNA) sprayed onto a crop is taken up by a fungal pathogen and silences an essential gene, is advancing as a biopesticide strategy. A recurring ambition is a broad-spectrum dsRNA: a single construct, directed at a conserved housekeeping gene, that controls several pathogens at once. Because silencing is driven by Dicer-processed small interfering RNAs (siRNAs) of roughly 21 nucleotides, the feasibility of this ambition depends on whether the target gene is conserved between pathogens at siRNA-length resolution, not merely at the level of overall gene identity. That resolution has not been mapped across the fungal plant-pathogen tree.

**Results:** We screened eleven canonical antifungal dsRNA target genes of *Fusarium graminearum* (three CYP51 sterol 14α-demethylases, six chitin synthases and two β-tubulins) for perfect 21-nucleotide identity against the transcriptomes of twelve fungal plant pathogens spanning four taxonomic classes (Sordariomycetes, Leotiomycetes, Dothideomycetes and Basidiomycota). Of 27,116 candidate siRNA start positions across the eleven genes, only 2,108 (7.8%) were conserved as a perfect 21-mer in even one other pathogen, 148 in at least three, twelve in at least seven, and none in eight or more. Conservation decayed steeply with phylogenetic distance: the congeneric *F. oxysporum* shared 1,209 sites, other Sordariomycetes 262–313, Leotiomycetes and Dothideomycetes 62–204, Basidiomycota 41–48, and the reduced-genome obligate biotroph *Blumeria graminis* just one. The β-tubulins carried the most broadly conserved individual sites (shared by up to seven pathogens), whereas CYP51C was the least conserved (shared by at most one). Even for the congener, perfect-match conservation covered only 4.5% of target positions and was concentrated in discrete windows rather than spread along the gene.

**Conclusions:** At the siRNA-length resolution that governs silencing, perfect-match conservation of canonical antifungal targets is scarce and strongly clade-limited. A perfect-match broad-spectrum dsRNA is therefore realistic only within a narrow clade, in practice around the genus level; genus- and class-spanning broad-spectrum SIGS cannot rely on perfect siRNA identity and would require mismatch-tolerant design or pathogen-specific constructs. The conserved windows mapped here identify where clade-level broad-spectrum constructs can be placed, and the gene ranking (β-tubulins most cross-targetable, CYP51C most pathogen-specific) provides a genome-guided basis for choosing antifungal dsRNA targets.

## Background

RNA interference (RNAi) has become a crop-protection technology. Host-induced gene silencing, in which a transgenic plant expresses RNAi constructs against a pathogen, was first shown for the powdery mildew fungus *Blumeria graminis* [1] and extended to *Fusarium graminearum* by silencing its three CYP51 sterol 14α-demethylase genes [2]. Spray-induced gene silencing (SIGS) removed the need for transgenesis: sprayed dsRNA is taken up by the fungus and processed by its own silencing machinery, controlling disease on treated tissue [3, 4]. The first sprayable dsRNA biopesticide against an insect pest has since been registered [5], and antifungal SIGS is the most mature of the remaining candidates [6].

A persistent goal in this field is a broad-spectrum dsRNA. If a single construct directed at a conserved, essential gene could suppress several pathogens, one product could replace several, which is attractive both economically and agronomically. The idea is not hypothetical. A dsRNA directed at conserved genes has been shown to protect plants against both *Sclerotinia sclerotiorum* and *Botrytis cinerea* [7], and the flagship antifungal construct CYP3RNA works by silencing all three *F. graminearum* CYP51 paralogues simultaneously through the sequence they share [4]. Longer constructs generate broader siRNA pools and greater cross-silencing potential [8]. The canonical antifungal target genes (sterol demethylases, chitin synthases and β-tubulins) are essential housekeeping genes, present as orthologues in every filamentous fungus and conserved under strong purifying selection, which makes them natural candidates for a construct intended to reach more than one pathogen.

Whether that ambition is genomically realistic depends on a quantity that overall gene identity does not capture. Silencing is triggered by small interfering RNAs (siRNAs), the roughly 21–25-nt products that Dicer-like enzymes cut from long dsRNA and load onto Argonaute, and Argonaute-directed cleavage requires extensive complementarity along the siRNA [9]. The operational unit of cross-pathogen targeting is therefore a contiguous stretch of about 21 nucleotides that is identical between the construct and the second pathogen, and regulatory bioinformatics for dsRNA biopesticides has converged on exactly this criterion, a perfect match of at least 21 nucleotides, as the primary sequence-based flag [10]. Two genes can be 80% identical overall and still share very few perfect 21-mers, because mismatches scattered along the alignment break the contiguous identical runs. The genome-level question for broad-spectrum design is thus not “how similar are these genes” but “how many perfect siRNA-length windows do they share, and where.” To our knowledge that question has not been answered systematically across the fungal plant-pathogen tree.

Here we map it. Using an exact k-mer method, we quantify perfect 21-nt conservation of the canonical antifungal target genes of *F. graminearum* against a panel of twelve fungal plant pathogens spanning four taxonomic classes, and we ask, for each gene, how many siRNA-length windows are shared with each pathogen and how that sharing scales with phylogenetic distance. A companion study used the same engine to screen these constructs against beneficial, non-target fungi and reported that perfect siRNA-length matches to biocontrol fungi are common [11]; here we turn the analysis on the pathogen axis, to define not the off-target hazard but the on-target scope for broad-spectrum design. The two analyses are complementary faces of a single property, the conservation of essential genes among related fungi, and we return to that link in the Discussion.

## Results

### An exact cross-pathogen conservation screen, validated against the design template

Eleven whole target transcripts of *F. graminearum* (CYP51A, CYP51B, CYP51C; six chitin synthases CHS1, CHS2, CHS3b, CHS4, CHS6, CHS7; and the β-tubulins TUB1 and TUB2) were decomposed into their constituent 21-mers on both strands, giving 27,116 candidate siRNA start positions. For each position we recorded in how many of the twelve pathogen transcriptomes an identical 21-mer occurs. Before any cross-pathogen result was read, the screen was validated against the design template itself: every one of the 27,116 positions was recovered in *F. graminearum*, a self-conservation of 100.0% for all eleven genes, confirming that the query construction and membership test behave as intended. The twelve pathogens (Table 1) span the Sordariomycetes (*F. oxysporum, Verticillium dahliae, Magnaporthe oryzae, Colletotrichum higginsianum, C. graminicola*), Leotiomycetes (*Botrytis cinerea, Sclerotinia sclerotiorum, Blumeria graminis*), Dothideomycetes (*Zymoseptoria tritici, Alternaria alternata*) and Basidiomycota (*Ustilago maydis, Puccinia graminis*).

**Table 1.** Plant-pathogen panel. Reference transcriptomes (cDNA) were retrieved from Ensembl Fungi release 63. *F. graminearum* is the design template from which the query genes were taken; the twelve screened pathogens are grouped by taxonomic class. (Perfect 21-nt sites = number of the 27,116 target positions conserved in that pathogen; see Fig. 1.)

| Class | Species | Assembly | Perfect 21-nt sites |
|---|---|---|---|
| Sordariomycetes | *Fusarium graminearum* PH-1 (template) | ASM24013v3 | 27,116 (self) |
| Sordariomycetes | *Fusarium oxysporum* | FO2 | 1,209 |
| Sordariomycetes | *Verticillium dahliae* | ASM15067v2 | 262 |
| Sordariomycetes | *Magnaporthe oryzae* (*Pyricularia oryzae*) | GCA900474545v2 | 313 |
| Sordariomycetes | *Colletotrichum higginsianum* | GCA_000313795.2 | 291 |
| Sordariomycetes | *Colletotrichum graminicola* | C_graminicola_M1_001_V1 | 264 |
| Leotiomycetes | *Botrytis cinerea* | ASM83294v1 | 71 |
| Leotiomycetes | *Sclerotinia sclerotiorum* | ASM14694v1 | 62 |
| Leotiomycetes | *Blumeria graminis* | EF2 | 1 |
| Dothideomycetes | *Zymoseptoria tritici* | MG2 | 113 |
| Dothideomycetes | *Alternaria alternata* | Altal1 | 204 |
| Basidiomycota | *Ustilago maydis* | Umaydis521_2.0 | 48 |
| Basidiomycota | *Puccinia graminis* | ASM14992v1 | 41 |

### Perfect siRNA-length conservation decays steeply with phylogenetic distance

Conservation of the target genes at siRNA resolution was low overall and fell sharply with distance from *F. graminearum* (Fig. 1). The congeneric *F. oxysporum* shared 1,209 perfect 21-nt sites, by far the most. The four other Sordariomycetes shared an order of magnitude fewer, between 262 (*V. dahliae*) and 313 (*M. oryzae*). Outside the Sordariomycetes the counts fell further: 62–71 in the Leotiomycetes *Botrytis* and *Sclerotinia*, 113 and 204 in the Dothideomycetes *Zymoseptoria* and *Alternaria*, and 41–48 in the Basidiomycota *Puccinia* and *Ustilago*. The obligate biotroph *Blumeria graminis*, whose genome is highly reduced and divergent, shared a single site across all eleven genes. The gradient is not a strict taxonomic ladder: *Alternaria* (Dothideomycetes) retained more perfect sites than the Leotiomycetes, reflecting lineage-specific rates of sequence divergence rather than rank alone. Even so, the dominant signal is unambiguous: perfect siRNA-length identity is a property that related fungi share and distant fungi do not.

**Fig. 1.**
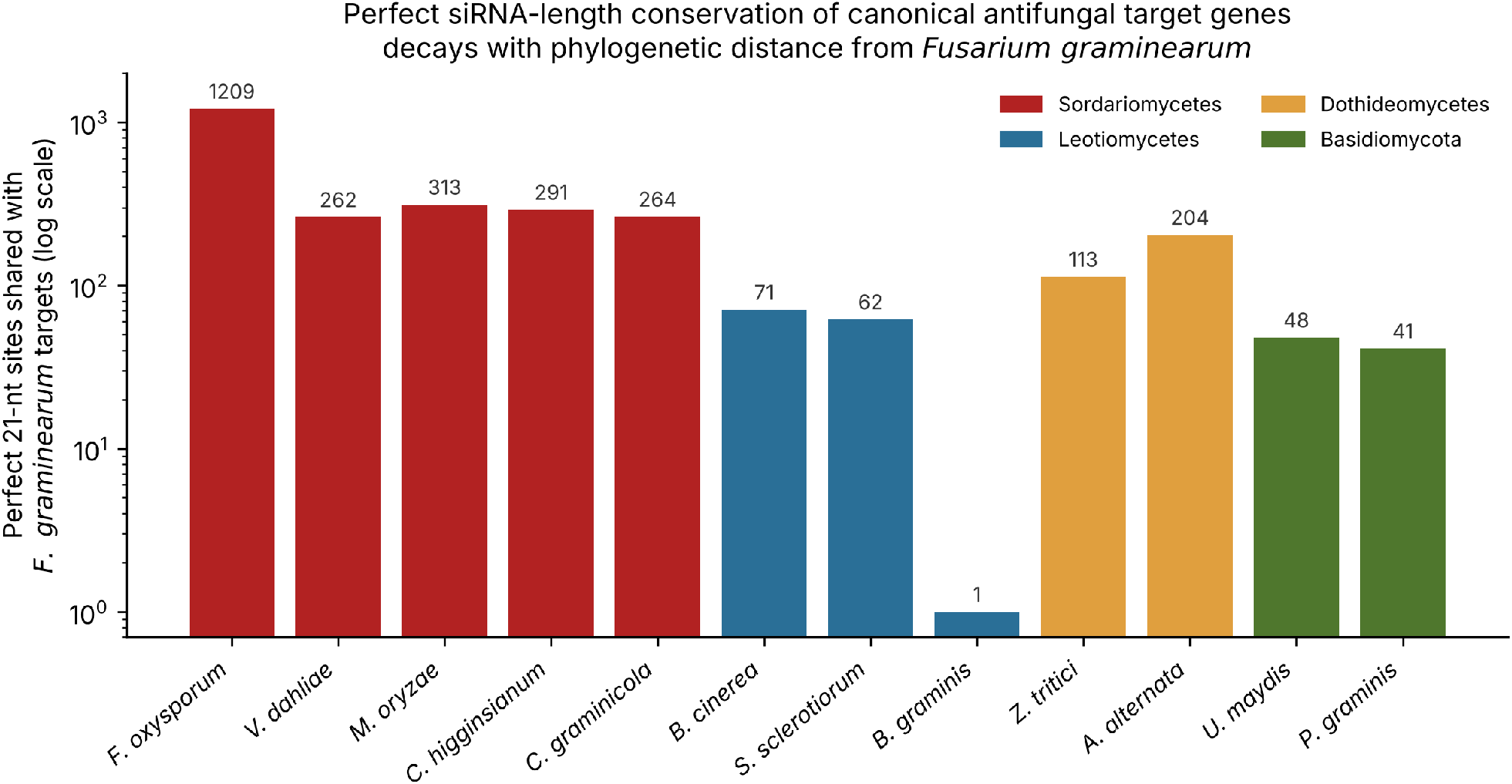
Perfect siRNA-length conservation of canonical antifungal target genes decays with phylogenetic distance from *Fusarium graminearum*. Bars show the number of the 27,116 target siRNA positions (eleven genes, both strands) that are conserved as a perfect 21-nt identity in each of the twelve pathogen transcriptomes, coloured by taxonomic class (log scale).

Across the whole panel the conserved fraction was small. Of the 27,116 target positions, only 2,108 (7.8%) were conserved as a perfect 21-mer in even one other pathogen; 440 (1.6%) in at least two, 148 in at least three, 53 in at least five, twelve in at least seven, and none in eight or more of the twelve pathogens (Fig. 3a). The single most broadly conserved siRNA site, in a β-tubulin, was identical in seven pathogens. Even the congener accounted for only 4.5% of all target positions, and, as shown below, those positions were concentrated in discrete windows rather than distributed along the genes.

### Conservation is gene- and lineage-specific, and the β-tubulins are the most cross-targetable

The per-gene, per-pathogen conservation matrix (Fig. 2; Table S1) shows that the shared sites are unevenly distributed across both axes. The β-tubulins were the most broadly conserved targets: TUB2 shared 104–145 perfect sites with each of the five Sordariomycetes, 30 with *Botrytis*, 103 with *Alternaria* and 24 even with the rust *Puccinia graminis*, and TUB1 showed the same pattern at lower magnitude. TUB1 and TUB2 also contained the only sites conserved in seven pathogens at once (Fig. 3b). At the opposite extreme, CYP51C was almost pathogen-specific: it shared nine sites with *F. oxysporum*, one with *C. higginsianum*, and none with any of the other ten pathogens, so that a construct placed in CYP51C would silence essentially *F. graminearum* alone. CYP51B and the chitin synthases CHS3b and CHS1 occupied an intermediate position, with substantial sharing among Sordariomycetes and little beyond.

**Fig. 2.**
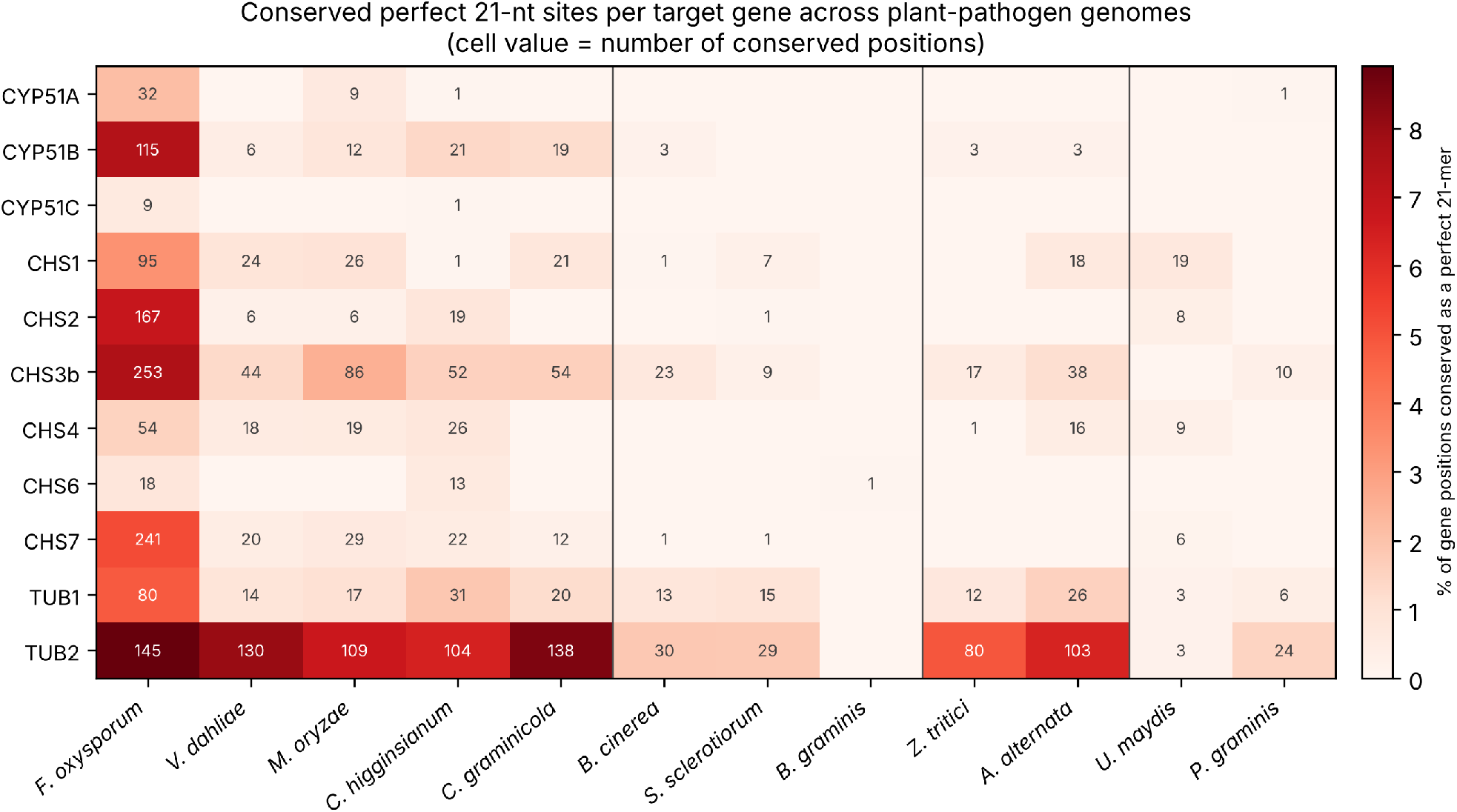
Conserved perfect 21-nt sites per target gene across the pathogen panel. Heat map of the number of conserved positions for each of the eleven target genes (rows) in each pathogen (columns), shaded by the percentage of the gene’s positions conserved; cell values give the absolute counts. Vertical lines separate taxonomic classes.

Expressed as a fraction of each gene, perfect-match conservation to the congener *F. oxysporum* ranged from 0.6% (CYP51C) and 0.8% (CHS6) to 7.4–8.9% (CYP51B, CHS3b, TUB2), confirming that even between congeners the cross-targetable sequence is a small, gene-dependent minority of the transcript.

### Broad-spectrum targeting is confined to a narrow clade

Taken together, the results bound where a perfect-match broad-spectrum dsRNA can work. Substantial cross-pathogen conservation, on the order of hundreds of shared siRNA sites, was confined to the genus *Fusarium*; across the rest of the Sordariomycetes it dropped to a few hundred sites concentrated in the most conserved genes, and beyond the class it collapsed to tens of sites or fewer. No single siRNA position was conserved across more than seven of the twelve pathogens, and the positions conserved across several pathogens were themselves a small, gene-biased set. A construct designed for perfect-match silencing can therefore be expected to reach a congener reliably and other members of the same class only partially and gene-dependently, and cannot be expected to reach pathogens in other classes through perfect identity at all.

## Discussion

The central result is a quantitative ceiling on broad-spectrum SIGS by perfect match. At the siRNA-length resolution that actually drives Argonaute-directed silencing, the canonical antifungal target genes share very little sequence between pathogens: under 8% of positions are conserved in even one other pathogen in our panel, and no position is conserved across more than seven of twelve. Conservation is steeply clade-limited, dropping roughly an order of magnitude from the congener to other Sordariomycetes and again beyond the class. The practical consequence is direct. A perfect-match dsRNA can be designed to reach a target pathogen and its close congeners, but a single construct cannot be expected to silence pathogens across genera or classes through perfect identity, because the shared siRNA-length windows that such silencing requires are largely absent. Broad-spectrum SIGS in the genome-wide sense, one spray for many unrelated pathogens, is not supported by the sequence data; broad-spectrum SIGS in the clade-restricted sense, one construct for a pathogen and its near relatives, is.

That distinction is actionable because the conserved sites are not scattered but concentrated, and because genes differ markedly in how far their conservation reaches. The β-tubulins carried the most cross-pathogen-conserved sites and the only sites shared by seven pathogens, which makes them the natural substrate for a clade-level broad-spectrum construct; the trade-off is that β-tubulin is also the benzimidazole fungicide target and, in the companion off-target analysis, the heaviest carrier of perfect matches to beneficial fungi [11], so its breadth cuts both ways. CYP51C sat at the other extreme, essentially pathogen-specific, and is therefore the gene of choice when the goal is to silence one pathogen and spare its relatives. CYP51B and the chitin synthases CHS3b and CHS1 offer intermediate, genus-level breadth. Choosing the gene, and then the window within it, is thus a genome-guided lever on the taxonomic range of a construct, available before any synthesis from public transcriptomes and a screen of the kind reported here, and complementary to the siRNA-design tools that predict dicing and guide-strand selection [12, 13].

The two faces of gene conservation meet here. The same purifying selection that preserves identical siRNA-length windows between a pathogen and its congeners also preserves them between a pathogen and the beneficial fungi that are its phylogenetic neighbours; the on-target scope for broad-spectrum design and the off-target hazard to related beneficials are produced by one property and scale together [11]. A gene or window chosen for maximal cross-pathogen reach will, for the same reason, carry the greatest risk of reaching non-target relatives, and a gene chosen for pathogen specificity minimises both. This coupling argues for treating target breadth as a design parameter set deliberately against both pathogen range and non-target safety, rather than maximised by default.

Several limitations bound the interpretation. First, we scored only perfect 21-nt identity. This is the stringent primary criterion of regulatory screening [10] and the most unambiguous unit of silencing, but it is a bound in two directions. It understates functional cross-reactivity, because siRNAs can also act through partial, seed-region complementarity [14], so two genes sharing few perfect 21-mers may still cross-silence through mismatched siRNAs; and it overstates the functional value of the perfect matches it does count, because dicing of a long dsRNA is non-uniform and not every 21-mer window is cut and loaded into Argonaute at comparable abundance [15, 16]. A perfect-match map therefore defines the high-confidence core of cross-pathogen targetability, not its full extent. Second, the analysis used one design template, the target genes of *F. graminearum*; a construct designed from a different pathogen’s orthologues would share its own, partly different, set of windows, and a full picture would screen each pathogen’s genes reciprocally. Third, the panel of twelve pathogens, although spanning four classes, samples the fungal tree sparsely and includes no oomycete or bacterial pathogen.

Fourth, conservation of sequence is necessary but not sufficient for silencing, which additionally requires environmental delivery, uptake, and a functional RNAi pathway in the recipient [3, 17]; the windows mapped here are candidate targets, not demonstrated ones. These limitations define the natural extensions: a mismatch-tolerant cross-pathogen screen, reciprocal designs from multiple templates, a wider panel, and experimental testing of candidate broad-spectrum windows.

## Conclusions

At the resolution that governs silencing, the canonical antifungal target genes are conserved between fungal plant pathogens far less than their housekeeping status suggests: perfect siRNA-length windows are scarce and sharply clade-limited, confined in practice to a target pathogen and its close relatives. Perfect-match broad-spectrum SIGS is therefore a clade-level, not a pan-fungal, strategy, and the gene and window chosen set its taxonomic reach. The conservation maps and gene ranking reported here, together with the recognition that cross-pathogen reach and off-target hazard to beneficial fungi are two expressions of the same conservation, provide a genome-guided basis for designing antifungal dsRNA constructs with a deliberately chosen spectrum.

## Methods

### Pathogen panel and target genes

Reference transcriptomes (cDNA) for *Fusarium graminearum* PH-1 and twelve fungal plant pathogens were obtained from Ensembl Fungi release 63 [18, 19] (Table 1). Eleven canonical antifungal dsRNA target genes were taken from the *F. graminearum* annotation and used as the design template: the sterol 14α-demethylases CYP51A, CYP51B and CYP51C (locus tags FGSG_04092, FGSG_01000, FGSG_11024) [2, 4]; six chitin synthases spanning classes I–VII (CHS1, FGSG_10327; CHS2, FGSG_02483; CHS3b, FGSG_10116; CHS4, FGSG_01272; CHS6, FGSG_01949; CHS7, FGSG_12039) [20, 21, 22]; and the β-tubulins TUB1 (FGSG_06611) and TUB2 (FGSG_09530, the benzimidazole target) [23]. Where a gene had multiple annotated transcripts the longest was used.

### Exact 21-nt conservation screen

The screen uses exact k-mer membership testing rather than alignment, implemented in Python with numpy and Biopython; the engine is the same as that described and deposited for the companion off-target study [11]. Each 21-mer was two-bit encoded into a 64-bit integer (A, C, G, T → 0, 1, 2, 3). For each target gene, the set of 21-mers on both strands (the forward 21-mer at each position and its reverse complement) was built, giving 27,116 siRNA start positions across the eleven genes; both strands were included because a dsRNA duplex is diced into siRNAs from each strand. Windows overlapping an ambiguous base were discarded. Each pathogen transcriptome was then streamed and every transcript 21-mer tested for membership in the target set; a target position was recorded as conserved in a pathogen if the pathogen transcriptome contained an identical 21-mer at that position on either strand. For each position we tallied the number of pathogens in which it was conserved (0–12), and for each gene–pathogen pair the number of conserved positions. As a positive control the template genome *F. graminearum* was screened against its own target set, which recovered 100.0% of positions for all eleven genes.

### Analysis

Per-pathogen totals (Fig. 1), the gene × pathogen conservation matrix (Fig. 2; Table S1), the distribution of positions conserved in at least k pathogens, and the per-gene maximum number of pathogens sharing a single site (Fig. 3) were computed from the per-position tallies. The complete analysis runs on a laptop from public data.

**Fig. 3.**
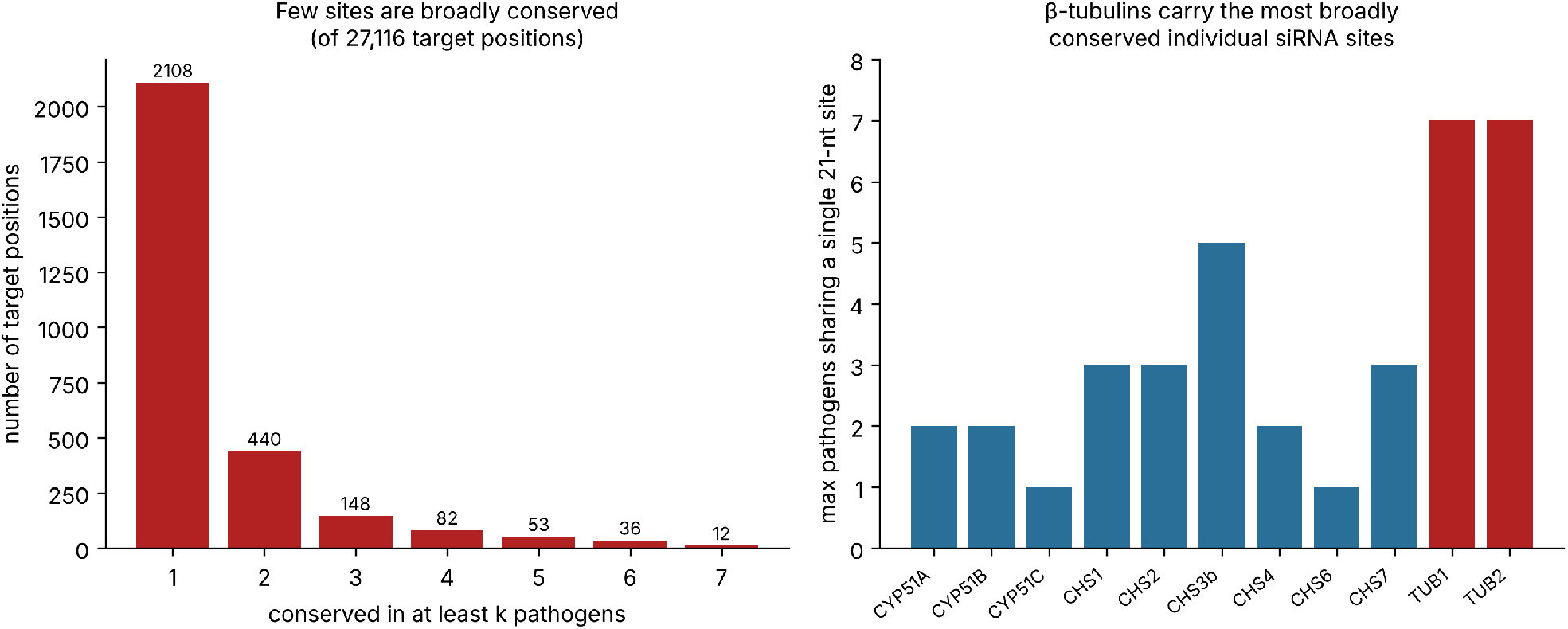
Broad-spectrum conservation is scarce. (a) Number of target positions conserved as a perfect 21-mer in at least k of the twelve pathogens; no position is conserved in more than seven. (b) The maximum number of pathogens sharing a single conserved site, per gene; the β-tubulins carry the most broadly conserved sites.

## Supporting information

Supplementary Code (analysis script)

## Declarations

### Availability of data and materials

All reference transcriptomes are publicly available from Ensembl Fungi release 63 [18, 19]. The k-mer screening engine and companion result files are openly archived at Harvard Dataverse (https://doi.org/10.7910/DVN/ZDLO7W, CC0 1.0) [24]; the cross-pathogen result files generated for this study are available from the corresponding author and will be deposited with the final version.

### Authors’ contributions

MW conceived the study, designed and performed the computational analysis, and drafted the manuscript. MAK and MAA contributed equally to this work, contributing to the design of the analysis, the interpretation of the results, and the revision of the manuscript. GT and HS contributed to the interpretation of the results and critically revised the manuscript; GT supervised the project. All authors read and approved the final manuscript.

## Acknowledgements

The authors thank colleagues at the University of Debrecen for helpful discussion.

## Use of generative AI

Generative AI (large language model) tools assisted with language editing, formatting and bibliographic checking. No AI tool was used to design the study, generate or analyse the data, or produce the scientific conclusions; the screen was performed with the authors’ own Python code. The authors reviewed all AI-assisted content and take full responsibility for it.

